# Influence of carbon-to-nitrogen ratio on the denitrifying activities of a methanol-fed, marine recirculating denitrification reactor, and the evolution of the bacteria community during the operating process

**DOI:** 10.1101/2025.05.02.651252

**Authors:** Livie Lestin, Richard Villemur

**Affiliations:** Institut national de la recherche scientifique (INRS), Centre Armand-Frappier Santé Biotechnologie, Laval, Province of Québec, Canada, H7V 1B7

**Keywords:** Denitrification, biofilm, carbon-to-nitrogen ratio, marine, methylotrophs, *Methylophaga*, nitrate, methanol

## Abstract

**Background:** Carbon-to-nitrogen ratio (C/N) is an essential parameter known to influence both the function and the activity of microbial communities in bioprocesses. The right balance between the contribution of each of these resources are essential for sustainable and cost-efficiency bioprocess. Here we aimed to assess the influence of C/N (here methanol and nitrate) on the performance of a recirculating denitrifying reactor for 31 weeks under marine conditions. We also monitored the evolution of its microbial community during the operating conditions.

**Methodology:** A 500-mL methanol-fed recirculating denitrification reactor operated under marine conditions and colonized by a naturally occurring multispecies denitrifying biofilm was subjected to eight different C/N over 31 weeks under anoxic conditions. We monitored several physico-chemical parameters (denitrifying activities, methanol consumption, CO_2_ production) throughout the operating conditions. Evolution of the bacterial community in the biofilm during the operating conditions was determined by 16S rRNA gene amplicon sequencing. Metatranscriptomes of representative conditions were performed to derive (1) the relative gene expression profiles of *Methylophaga nitratireducenticrescens* strain GP59, the main denitrifier, and (2) the functional diversity of the biofilm.

**Results:** Changes in C/N did not impact the denitrifying activities of the recirculating reactor but did impact the carbon dynamics. Throughout the operating time, nitrite and N_2_ O appeared transiently, and ammonium was not observed. The bacterial community in the reactor increased in diversity as the biofilm aged, especially in heterotrophic bacteria, at the expense of methylotrophic bacteria. The functional diversity suggests that heterotrophs could use formaldehyde, which may have been released by methylotrophs in the biofilm, as energy and carbon source. The relative expression profiles of strain GP59 in the biofilm is distinct of those of GP59 planktonic pure cultures, and that the expression of several riboswitches and *xoxF* would be involved in such differences.

**Conclusions:** When the biofilm community is well established in the reactor, it can sustain changes in C/N with limited impact on the denitrification performance. Increase in the proportion of heterotrophs would allow the reactor to be more flexible for carbon sources. Such knowledge can be useful in the optimisation of denitrification process.

## 1 Introduction

Nitrate (NO_3_^−^) can be a threat to water ecosystems (Rouse *et al*., 1999, Camargo *et al*., 2005), which is specifically true for close-circuit ecosystems such as aquarium, where NO_3_^−^ can reach high level (>200 ppm) (Grguric & Coston, 1998), and water replacement has to be performed. However, water replacement can be costly and stressful to the ecosystem (Martins *et al*., 2010). The main water treatment of these aquariums does not remove efficiently NO_3_^−^, and denitrification system is added to remediate to the problem (Müller-Belecke *et al*., 2013).

Denitrification is a microbial process where N oxides serve as terminal electron acceptors instead of oxygen (O_2_) for energy production when O_2_ depletion occurs, leading to the production of gaseous nitrogen (N_2_). Four sequential reactions are required where NO_3_^−^ is reduced to N_2_, via nitrite (NO_2_^−^), nitric oxide (NO) and nitrous oxide (N_2_ O), and each of these reactions is catalyzed by different enzymes, namely NO_3_^−^ reductases (Nar and Nap), NO_2_^−^ reductases (NirS and NirK), NO reductases (Nor) and N_2_ O reductases (Nos) (Philippot & Hojberg, 1999, Richardson *et al*., 2001, Kraft *et al*., 2011). Different denitrification processes and strategies have been developed over decades, each in response to specific characteristics of infrastructure and water to treat (Ni *et al*., 2017).

Bioprocesses are composed of an amalgam of microorganisms arranged in a microbial community involved in achieving the biochemical reactions necessary for transformations of the pollutants of interest. The microbial community is in general impacted in the composition of its populations by the operating conditions of the bioprocess, such as, among others, the water type (e.g. freshwater, seawater), the type of molecules to transform, the nutrients provided for electron donors, the carbon source, co-factors, and electron acceptors like O_2_ under oxic conditions or N oxides (e.g. NO_3_^−^) under anoxic conditions. For instances, methanol-fed denitrification processes are composed mostly of methylotrophic bacteria (used C1 carbon as carbon source), which is different from acetate-fed processes, where mainly heterotrophic bacteria composed their microbial communities (Lu *et al*., 2014). Heterotrophic/methylotrophic denitrification processes (as opposed to autotrophic denitrification) need a readily carbon source to achieve denitrification (Ni *et al*., 2017, Fu *et al*., 2022, Brozincevic *et al*., 2024). As readily available carbon can be low in water system to treat, external carbon has to be added such as acetate, ethanol and methanol. Methanol has the advantage to be not costly and available. The methanol-fed processes produce low biomass; thus, a higher proportion of carbon is used as electron donors for the denitrification (Ni *et al*., 2017). However, a methylotrophic community has to establish in these systems, which may pose a problem of lagging time, and are not necessarily flexible as only C1 carbon can be used, contrary to the heterotrophic systems.

Among factor that can affect denitrification performance is the right level of carbon to feed the bioprocess for optimal activities. The carbon-to-nitrogen ratio (C/N) is an essential parameter, known to influence both the function and the activity of microbial communities in bioprocesses (Lu *et al*., 2014). Under denitrifying conditions (anoxic), NO_3_^−^ (and N oxides) replaces O_2_ as terminal electron acceptors for respiration, which is intimately linked to carbon assimilation. In methanol-fed denitrification processes, methylotrophic bacteria use methanol as source of carbon and energy, where it is first converted to formaldehyde by the methanol dehydrogenase with transfer of electrons to cytochromes and subsequently to the respiratory chain. Formaldehyde can be either assimilated in the biomass, or dissimilated in formate and CO_2_ by enzymatic reactions that generate NADH (Anthony, 1982).

Adjustments of C/N are important to avoid, for instances, an inadequate methanol dosage when operating a denitrification process. Non-metabolized methanol (over dosage) could pollute the effluent and potentially harm the ecosystem. Also, non optimal dosage can result in incomplete denitrification, releasing toxic molecule such as N_2_ O, NO_2_^−^, or H_2_ S especially in marine system (high level of sulfate), or induce the dissimilatory NO_3_^−^ reduction to ammonium (DNRA) pathway (Her & Huang, 1995, Cattaneo *et al*., 2003, Yang *et al*., 2012, Mohan *et al*., 2016). Therefore, the right balance between the contribution of each of these resources are essential for sustainable and cost-efficiency denitrification process.

The stoichiometry of the denitrification reaction with methanol, 5CH_3_ OH + 6NO_3_^−^ => 3N_2_ + 5CO_2_ +7H_2_ O + 6OH^−^, requires in theory 0.714 g-C methanol/g-N NO_3_^−^ (C/N of 0.7) to achieve complete denitrification. However, as denitrification is a microbial process, where methanol and NO_3_^−^ are also involved in assimilation for growth and cell maintenance, and that nitrite and O_2_ can be present in the treating water, higher level of methanol is needed (McCarty *et al*., 1969, Tchobanoglous *et al*., 2003). For instances, Yin and Guo (2022) and Her and Huang (1995) tested different C/N on sequencing batch reactors (SBR) with sludge acclimated for denitrification. They showed that with C/N of 1.1 and 0.9, respectively, the reactors achieved near 100% nitrogen removal.

We have been studying for the last two decades the microbial ecology of a denitrifying biofilm that developed in a fluidized-bed, methanol-fed denitrification system operating in continuous mode for treating the seawater tank at the Montreal Biodome natural museum. This biofilm is composed of an autochthonous methylotrophic community of at least 15 bacterial species (Labbé *et al*., 2003, Labbé *et al*., 2007, Laurin *et al*., 2008), from which several strains were isolated; among others, the ones involved in denitrifying activities are *Hyphomicrobium nitrativorans* strain NL23, and strain JAM1 and strain GP59 of the species *Methylophaga nitratireducenticrescens* (Auclair *et al*., 2010, Martineau *et al*., 2013, Villeneuve *et al*., 2013, Geoffroy *et al*., 2018). We use this denitrifying biofilm as a model to study how microbial communities of bioprocesses evolve in response to changes during bioprocess operations. For instance, we cultivated the biofilm at laboratory-scale (vials) to assess its denitrification performance towards physico-chemical changes such as the type of medium, the temperature, pH, and the concentrations of NaCl, methanol and NO_3_^−^, and to further assess the impact of these changes on the microbial community (Payette *et al*., 2019, Villemur *et al*., 2019). Our results have demonstrated that, when the microbial community is well-established, it remains stable and can sustain denitrifying activities despite of all these changes. However, this approach involved static, batch culture assays and artificial seawater, which differed substantially from the continuous operating conditions and the seawater medium of the original denitrification system. To cope with this problem, we developed a 500-mL recirculating bioreactor to mimic as much as possible at the laboratory scale the continuous operating conditions. We succeeded in developing biofilm in this reactor by co-culturing *M. nitratireducenticrescens* strain JAM1 and *H. nitrativorans* strain NL23, with sustainable denitrifying activities (Cucaita *et al*., 2021).

Although numerous studies assessed the effect of C/N on diverse types of denitrification systems, relatively few studies have examined the influence of C/N changes on these systems operated under methylotrophic and marine conditions. We aimed in the present work to assess the influence of C/N (here methanol and NO_3_^−^) on the performance of the denitrifying methylotrophic biofilm process operated under marine conditions. Our recirculating reactor was inoculated with the biofilm taken from the Biodome denitrification reactor, in which fresh biofilm developed on supports. The reactor was then running for 31 weeks and operating sequentially under with eight different C/N. The impact of these C/N was assessed on several physico-chemical parameters measured during the operating conditions. We also monitored the evolution of bacterial community to assess the stability of the bacterial community during the operating conditions. Using metatranscriptome approach, we derived the functional diversity of the biofilm, which indicated the potential metabolic pathways occurring in the active microbial populations. Finally, the relative gene expression profiles of strain GP59 in the recirculating reactor were compared to those from our previous works, planktonic cultures for instances (Payette *et al*., 2019, Villemur *et al*., 2019, Lestin & Villemur, 2024), to broaden our study to assess the impact of the physiology and the operating modes on these expression profiles. Better understanding how C/N can impact the denitrification performance under methylotrophic, marine conditions, and how the microbial populations response to these changes will provide useful information for optimizing these denitrification processes.

## 2 Material and Methods

### 2.1 Acclimation of denitrifying biomass in recirculating reactor

The medium was the commercial seawater Instant Ocean (IO) medium (Aquarium systems; Mentor, OH, USA), the same used in the original denitrification process. The IO medium (30 g/L) was autoclaved, and 1 mL/L of sterilized trace element solution (FeSO_4_ •7H_2_ O 0.9 g/L; CuSO_4_ •5H_2_ O 0.03 g/L; MnCl_2_ •4H_2_ O 0.234 g/L; Na_2_ MoO_4_ •2H_2_ O 0.363 g/L) was added.

The denitrifying biomass was provided in the form of biofilm, immobilized on polyethylene supports of the Bioflow 9-type (1.2 mm diameter) that had developed in the denitrification reactor at the Montreal Biodome, but was preserved in 20% glycerol (in IO medium) at −20°C (Payette *et al*., 2019). Frozen supports were thawed and submerged for 5 days in the IO medium at 4°;C, supplemented with 0.3% methanol. The biofilm was also scraped from the supports and inoculated in the same medium. The biomass free or attached to the supports was transferred to the reactor with seventy empty Bioflow 9-type supports. These empty supports were previously washed in a 35% hydrochloric acid bath overnight and then rinsed a minimum of three times with demineralized water.

The reactor consisted of a glass vessel with an airtight 550-mL working volume, installed in a glove box continuously purged with nitrogen gas and equipped with an oximeter to confirm the absence of O_2_. A 500-mL volume of culture medium supplemented with 0.3% methanol and 21.4 mM NaNO_3_ (300 mg-N/L) (C/N of 3) was added to the reactor. A 4-L tank filled with the same medium was appended to the circuit during the acclimation period. The medium was recirculated with a peristaltic pump at 20 mL/min, and the reactor was run at room temperature (ca. 22 °C). The excessive volume of gas produced due to the denitrifying activities was evacuated in a graduated bottle inverted in water and connected to the reactor. Gas production was recorded by water displacement. All equipment, including culture medium, was sterilized before the start of the experiment; however, the operation of the reactor was not carried out under sterile conditions, although it was operated in closed environment of the glove box. The culture medium was replaced regularly as soon as NO_3_^−^ was exhausted. When biofilm was apparent on the empty supports, supports with the original biofilm were removed. When denitrifying activities was stable, the reactor was emptied, and half supports were set aside and stored at 4°C in IO medium for future used. From this, the C/N assays were carried out with appropriate NO_3_^−^ and methanol concentrations.

### 2.2 C/N assays

The reactor was run sequentially under eight different C/N ranging from 1.5 to 7.5 according to Table 1, by adjusting the concentration of either NO_3_^−^ or methanol. The reactor was operated with medium recirculated at 20 mL/min (not with the 4L tank) in sequential batch mode, according to a cycle of three phases: a filling phase, a reaction phase that lasted until complete NO_3_^−^ reduction, and an emptying phase. The eight conditions were applied twice, which we refer as Period 1 (P1) and Period 2 (P2). The reactor ran from November 13, 2019, to June 14, 2020 (31 weeks). During Condition 6 (Period 2), the running operation was abruptly interrupted because of the Covid-19 crisis, which lasted 68 days, during which the supports were transferred at 4°C in IO medium containing NO_3_^−^ and methanol. During P1, the operating time for each condition ranged from 96 to 288 h with 1 to 3 medium changes. During P2, the same conditions were reapplied over a shorter interval for Conditions 1 to 5 (48 to 72h) as each condition was running with no medium changes. For Conditions 6 to 8, the reactor was running during much longer time (120 to 240 h) because of the decrease in the denitrification performance probably due to the storing period (Covid crisis).

**Table 1.** Operating conditions of the reactor.

|  | NO<sub>3</sub><sup>-</sup><br>mM | Methanol<br>v/v, % | C/N<br>ratio | P1<br>h | P2<br>h |
| --- | --- | --- | --- | --- | --- |
| C1 | 10.7 | 0.075 | 1.5 | 96 (2) | 48 (1) |
| C2 | 10.7 | 0.15 | 3.0 | 192 (4) | 48 (1) |
| C3 | 21.4 | 0.15 | 1.5 | 144 (2) | 72 (1) |
| C4 | 21.4 | 0.3 | 3.0 | 144 (2) | 72 (1) |
| C5 | 21.4 | 0.45 | 4.5 | 144 (2) | 72 (1) |
| C6 | 21.4 | 0.75 | 7.5 | 144 (2) | 240 (2) |
| C7 | 32.1 | 0.45 | 3.0 | 288 (2) | 120 (1) |
| C8 | 42.8 | 0.6 | 3.0 | 192 (2) | 192 (1) |

The reactor was first run sequentially from Condition 1 (C1) to Condition 8 (C8) and referred as Period 1 (P1). The reactor was then operated again under these eight conditions and referred as Period 2 (P2). During P2, Condition 6 (C6-2), the reactor operation was stopped abruptly, and the reactor was set aside at 4°C for several weeks because of the Covid-19 crisis. After this unfortunate situation, the reactor operation was resumed under Condition 6. The columns P1 and P2 refer to the operating time in hours and number under parentheses is the number of times the reactor was operated with fresh medium.

Liquid samples (one mL) were taken at sampling ports and centrifuged; the supernatant was used to determine the concentrations NO_3_^−^, NO_2_^−^, ammonium and methanol. The pellets of some of these samples were kept for DNA or protein (or both) extractions. A 5-mL volume of gas was also taken from the head space to measure N_2_ O and CO_2_.

At the end of each condition at the time of the emptying phase, three supports were taken to determine the protein content of the biofilm and to extract the nucleic acids (DNA and RNA). These supports were replaced by three others from the supports that were set aside previously, in order to keep a fixed number of supports. At the emptying phase, the medium was also recovered to collect the suspended biomass and determine its protein content. The biomass collected to extract the RNAs was scraped from supports under the anoxic environment of the glove box and immersed in the extraction buffer (50 mM Tris-HCl, 100 mM EDTA, 150 mM NaCl, pH 8.0) and water-saturated phenol pH 4.3 (v/v). The sample tubes were then immediately frozen in liquid nitrogen and stored at −70°;C. For DNA extraction and total protein assays, the biomass from the supports was scraped and then frozen at −20°;C in 5 mL of sterile culture medium. To recover the biomass in suspension, 300 mL of the culture medium was centrifuged at 16 000 x g and the pellet was mixed with 5 mL of sterile medium, and stored at −20°;C.

### 2.3 Analytic methods

Measurements of the concentrations of NO_3_^−^, NO_2_^−^ and protein content, and the determination of the specific NO_3_^−^ reduction rates and the specific NO_x_ (NO_3_^−^ + NO_2_^−^) rates were carried out according to Cucaita et al. (2021). N_2_ O and CO_2_ concentrations (in ppmv) in the headspace were determined by gas chromatography. Headspace samples (10 mL) were collected using a Pressure Lok gastight glass syringe (VICI Precision Sampling Inc., Baton Rouge, LA, USA) and were injected through the injection port of a gas chromatograph equipped with a thermal conductivity detector and electron-capture detector (7890B seriesGCCustom, SP1 option 7890-0504/0537; Agilent Technologies). The amount of N_2_ O (in µmole) in the reactor was calculated as described in Mauffrey et al. (2017). The NO_2_^−^ and N_2_ O showed transitory dynamics with appearance that peaked after a certain time, then reduced completely. To quantify these dynamics, we calculated the area under the curve of the amount of NO_2_^−^ or N_2_ O by the time (mM-h) they appeared and then disappeared. The CO_2_ production rates were calculated from the linear portion of CO_2_ produced by the time, which occurred mainly during the first 5-15 h. This period corresponded when NO_3_^−^ reduction was occurring.

Methanol was measured by gas chromatography coupled with a flame ionization detector (GC-FID). Nitrogen gas was used as a carrier gas. The culture medium samples were supplemented with 5% isopropanol as an internal standard. The standard range included solutions of 0.01 to 0.2% methanol. Total amount of methanol consumed was determined at the end of the reaction phase. The methanol consumption rates followed a first order dynamics, where a k factor was calculated as [ln(C_t_ /C_0_)]/time with C_t_ methanol concentration at the sampling time and C_0_ the initial methanol concentration. Ammonium concentration was measured by colorimetric method as described by Mulvaney (1996).

### 2.4 PCR-DGGE and 16S rRNA amplicon sequencing

DNA extractions of biomass on support were performed as described (Geoffroy *et al*., 2018). The PCR amplifications of the V3 region of the 16S rRNA genes for denaturing gradient gel electrophoresis (DGGE) experiments were performed as described (Lafortune *et al*., 2009) with the 341f and 534r primers.

The microbial composition of the biofilm was determined by amplicon sequencing of the 16S rRNA V3-V4 regions with the S-D-Bact-0341-b-S-17 (5’ <u>TCGTCGGCAGCGTCAGATGTGTATAAGAGACAG</u>-CCTACGGGNGGCWGCAG 3’) and S-D-Bact-0785-a-A-21 (<u>GTCTCGTGGGCTCGGAGATGTGTATAAG</u> <u>AGACAG</u>-GACTACHVGGGTATCTAATCC) primers (Klindworth *et al*., 2013) linked with Illumina sequences (underlined sequences). The amplifications were carried out in 25-µL volume containing the commercial Taq polymerase buffer, 0.25 mM MgSO_4_, 0.2 µg BSA, 200 nM of each primer, 0.5-unit AccuPrime™ Taq DNA Polymerase (Invitrogen, Carlsbad, CA, USA) and 50 ng DNA extract. The reactions were run at 94°;C 5 min, 30 cycles at 94°;C 30 sec, 55°;C 30 sec, and 68°;C 30 sec, and finally at 68°;C 10 min. Residual primers and primer dimers were removed with the AMPure XP magnetic beads kit according to the manufacturer (Beckman Coulter, USA). The amplicons were then amplified a second time in which indexes were added. The amplifications were carried out reaction in 25-µL volume containing the Taq polymerase buffer, 0.25 mM MgSO_4_, 0.2 µg BSA, 400 nM index primers, 0.5-unit AccuPrime™ Taq DNA Polymerase (Invitrogen, Carlsbad, CA, USA) and 5 µL amplicon. The reactions were run at 94°;C 3 min, 8 cycles at 94°;C 30 sec, 55°;C 30 sec, and 68°;C 30 sec, and finally at 68°;C 5 min. Residual primers and primer dimers were once again removed with AMPure XP magnetic beads kit, and the purified amplicons were quantified with the Quant-iT PicoGreen dsDNA Assay kit (Invitrogen). The amplicons were sent for sequencing by Illumina MiSeq PE250 (250 bp paired-end sequencing reactions) at the Centre d’expertise et de services Génome Québec (Montréal, QC, Canada). Sequencing reads were processed by the dada2 pipeline at the Galaxy server (https://usegalaxy.org/) to generate ASVs (amplicon sequence variants). The taxonomic assignment was carried out at the Silva web site (https://www.arb-silva.de/). Number of reads ranged from 27 268 to 33 031 for the recirculating reactor samples, and 423 208 reads from the original biofilm sample. Rarefaction analyses were performed with the Analytic Rarefaction v1.3 software downloaded from the UGA Stratigraphy Lab web site (stratigrafia.org/) with increments of 10. All samples showed rarefaction saturation. ASVs affiliated to the same lineage were grouped together, in order to establish the proportions of each taxon in the samples. Raw sequencing data were deposited in Sequence Read Archive (SRA) at the National Center for Biotechnology Information (NCBI: https://www.ncbi.nlm.nih.gov/).

### 2.5 Metatranscriptomes

The RNA extraction was previously described by Mauffrey et al. (2015). RNA quality was verified by agarose gel electrophoresis and by spectrophotometry (Nanodrop) with ratio *OD*_*260*_ /*OD*_*280*_ >1.8. The RNA samples were sent for sequencing using the Illumina Method (NovaSeq 6000 S4 PE100). Library preparation and sequencing were performed by the Centre d’expertise et de services Génome Québec (Montréal, QC, Canada). Ribosomal RNA were depleted using the Ribo-ZeroTM rRNA Removal Kits (Meta-Bacteria; Epicentre, Madison, WI, USA). RNAseq reads were deposited in SRA. Raw sequencing reads were trimmed using fastq_quality_filter (1.0.2) to remove low quality reads (score < 20) (Gordon & Hannon, 2010), and the paired reads were then merged.

### 2.6 Relative gene expression profiles of strain GP59

The paired reads were aligned to the genome and plasmids of *M. nitratireducenticrescens* strain GP59 (GenBank accession number CP021973.1, CP021974.1, CP021975.1) using Bowtie2 (v 2.5.0) (Langmead & Salzberg, 2012) and annotated with Bedtools (v 2.30.0) (Quinlan & Hall, 2010). Genes that were significantly differentially expressed were identified by EdgeR (v 3.36.0) (Robinson *et al*., 2010) with trimmed mean of M values (TMM) method to normalize library sizes (robust=TRUE; P-value adjusted threshold=0.05; P-value adjusted method=Benjamini and Hochberg). All these analyses were performed on the Galaxy server (https://usegalaxy.org/). Genes were considered differentially expressed when the false discovery rate (FDR) was ≤ 0.05.

### 2.7 Analysis of the unaligned RNA reads

The paired reads were aligned using Bowtie2 to a concatenated sequence consisting of the three reference genomes (*M. nitratireducenticrescens* strain JAM1 [GenBank accession number CP003390.3] + strain GP59 + *H. nitrativorans* strain NL23 [CP006912]) and the two GP59 plasmids. The reads that did not align were *de novo* assembled by Trinity v. 2.4.0 (Grabherr *et al*., 2011) at the Galaxy server. Estimation of the relative transcript abundance of the *de novo* assembled sequences was performed by RSEM (Galaxy server) (Li & Dewey, 2011) and expressed as Transcript per million by RSEM [TPM-RSEM]. The resulting assembled sequences (contigs) were annotated at the Joint Genomic Institute (https://img.jgi.doe.gov/cgi-bin/m/main.cgi) to find open reading frames (ORFs) with their putative function and affiliation.

Contigs containing ORF (contig-ORF) and showing affiliation to a microbial lineage were grouped by taxon and their TPM-RSEM were summed. The proportion of each taxon per biofilm sample was then derived. For each identified taxon, the KO enzyme number from deduced function of gene products were retrieved and used to do reconstruction metabolic pathway by KEGG mapper-reconstruct analysis (https://www.genome.jp/kegg/ mapper/reconstruct.html). If the identified taxa were also represented by a corresponding species in KEGG database, the metabolic pathways were examined. The carbon assimilation and dissimilation pathways were derived from KEGG (https://www.genome.jp/kegg/pathway.html) and MetaCyc (https://metacyc.org) databases.

## 3 Results

### 3.1 Acclimation of the denitrifying biomass

The denitrifying biofilm taken from the denitrification reactor at the Montreal Biodome was used to inoculate the recirculating reactor for the colonization of new supports. We refer to this biofilm as the *Original Biofilm* to distinguish it from the biofilm that developed in the recirculating reactor. The reactor was operated under sequential-batch conditions with the IO medium, the same used by the Biodome, and supplemented with 21 mM NO_3_^−^ and 0.3% methanol (C/N of 3). After 37 days, stable denitrifying activities (NO_3_^−^ and NO_2_^−^ reduction) occurred in the reactor, and fresh biofilm was visually observed on the supports (Fig. 1). We refer to this biofilm as the *RR biofilm* (RR for recirculating reactor). The C/N assays were then carried out. Day 0 (D0) was set after the 37-day acclimation period.

**Figure 1:**
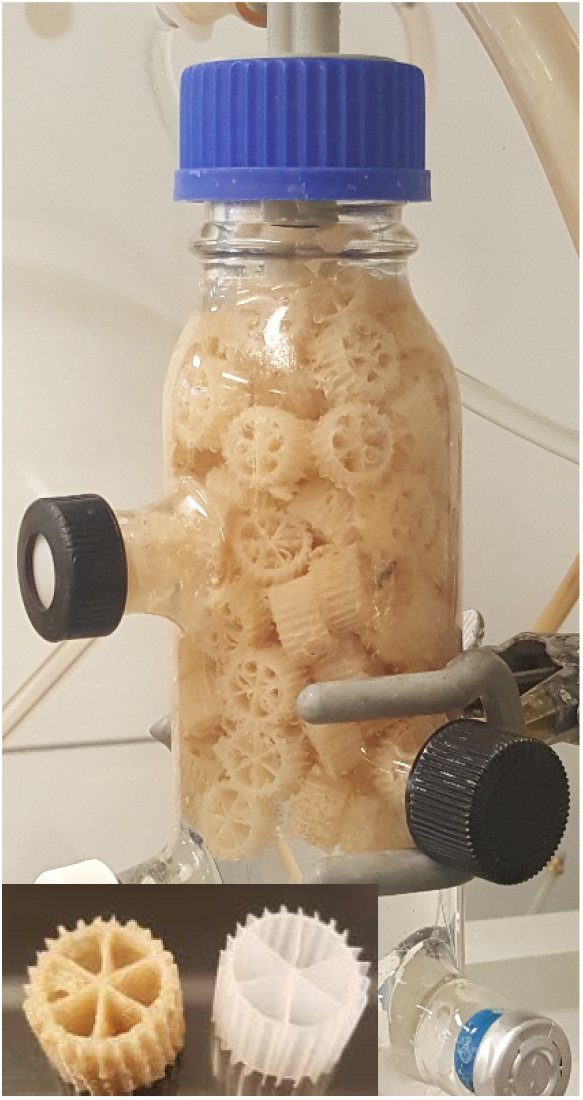
Biofilm on the reactor supports. Panel A. Recirculating reactor. Panel B. Left: support with biofilm. Right: virgin support

### 3.2 C/N assays

The reactor with fully colonized supports by the denitrifying biomass was run sequentially under eight different C/N (1.5 to 7.5) (Table 1). These conditions were performed twice (P1 and P2) with the same biomass to assess the reproductivity of the results. The reactor ran for 31 weeks during these two periods, where several physico-chemical parameters were measured (Fig. 2). pH was stable (7.6-8.0), and protein content on the supports (35), representing the total microbial biomass, ranged from 10 to 25 mg. No ammonium was detected in all conditions.

**Figure 2.**
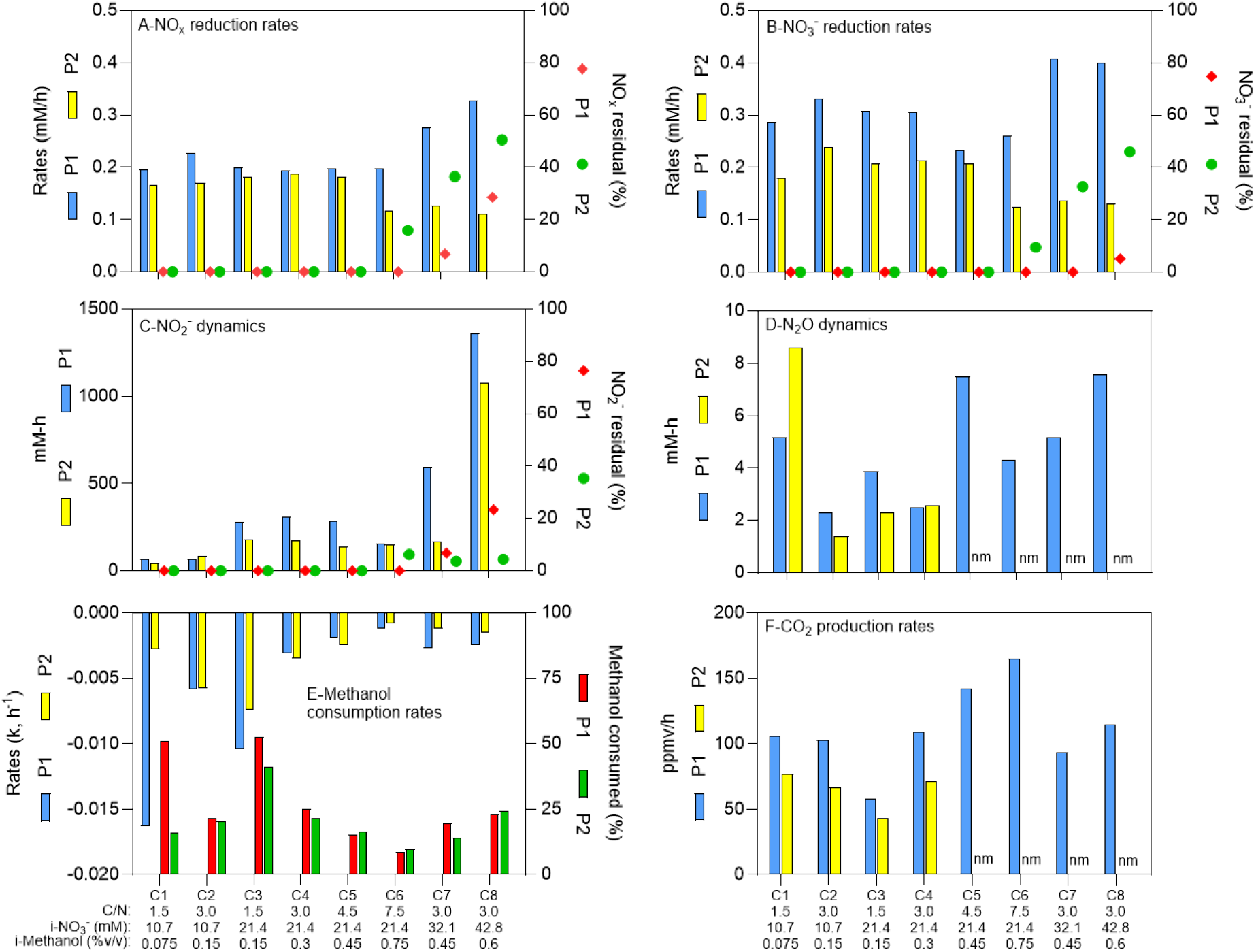
Physico-chemical measurements during the different operating conditions. See Table 1 for conditions’ nomenclature. Residuals: % of NO_x_, NO_3_^−^ and NO_2_^−^ from the initial methanol concentrations (i-NO_3_^−^) that remained in the reactor after four operating days. Methanol consumption rates followed first order dynamics. Methanol consumed: % of methanol consumed at the end of the period. nm: not measured.

NO_3_^−^ reduction was occurring with transient NO_2_^−^ and N_2_ O accumulation (Fig. 2C and D). In average, the NOx (NO_3_^−^ + NO_2_^−^) reduction rates (Fig. 2A) were 77% of NO_3_^−^ reduction rates (Fig. 2B), because of the transient NO_2_^−^ accumulation (Fig. 2C). Although we operated twice the reactor under the eight C/N ratios, we encountered an important obstacle, the Covid-19 crisis, where the reactor had to be stopped abruptly during P2-Condition 6, and stored in cold room for several weeks that negatively impacted the denitrification performance (Fig. 2). If we assume the NOx reduction rates representing the denitrification rates (NO and N_2_ O reduction has minimal influence on these rates), the performance of the reactor ranged from 0.166 to 0.328 mM h^-1^ if we exclude post-Covid data. These rates averaged 0.119 mM h^-1^ in P2-Conditions 6 to 8. We estimated the residuals of NOx, NO_3_^−^ and NO_2_^−^ after four operating days (Fig. 2A-C). We observed these residuals in P1 only under Conditions 7 and 8, with 28.5% residual NOx in Condition 8. In P2, Conditions 6 to 8 showed higher levels of residuals because of the lower reactor performance with 50.6% residual NOx in Condition 8.

The peak of N_2_ O concentrations appeared mostly in the first 10h to 20h of operating conditions and ranged from 0.04 mM to 0.4 mM. No residual N_2_ O was found at the end of each condition. The amount of methanol consumed varied from 0.031% to 0.147% (v/v), which corresponded between 9% to 53% of the initial methanol concentrations (i-Methanol) (Fig. 2E). Methanol consumption rates (Fig. 2E) followed first order dynamics, whereas the CO2 production rates (Fig. 2F) were at their highest during the first 5-15 h.

Correlation analyses were first performed between (i) the i-Methanol, (ii) the initial NO_3_^−^ concentrations (i-NO_3_^−^), (iii) the C/N ratio applied to the reactor, and the physico-chemical parameters measured during the operating conditions (Table 2). It revealed that changes in C/N did not correlate with the denitrification dynamics (NO_3_^−^ reduction rates, the NOx reduction rates, the NO_2_^−^ and N_2_ O dynamics; p>0.05). However, the methanol consumption rates, and the CO2 production rates did correlate with the C/N ratios (0.05<p<0.001; Table 2). Correlation (p<0.05) occurred between the i-NO_3_^−^ and (i) the NO_3_^−^ reduction rates, (ii) the NOx reduction rates and (iii) the NO_2_^−^ dynamics (Table 2). The i-Methanol correlated with the methanol consumption rates and the CO2 production rates (0.05<p<0.001; Table 2). Correlation analyses were also performed between the respective measured parameters (Table 2). As expected, strong correlation (p<0.0001) was observed between the NO_3_^−^ reduction rates and the NOx reduction rates (Table 2). The N_2_ O dynamics did not correlate with any conditions.

**Table 2.** Pearson correlation coefficients between measured parameters.

| | $\text{NO}_3^-$<br>reduction<br>rates | $\text{NO}_x$<br>reduction<br>rates | $\text{NO}_2^-$<br>dynamics | $\text{N}_2\text{O}$<br>dynamics | Methanol<br>consumption<br>rates | $\text{CO}_2$<br>production<br>rates |
| --- | --- | --- | --- | --- | --- | --- |
| i-$\text{NO}_3^-$ | 0.0312 | 0.0009 | <0.0001 | ns | ns | ns |
| i-Methanol | ns | ns | ns | ns | 0.0154 | 0.0065 |
| C/N | ns | ns | ns | ns | 0.031 | 0.0015 |
| $\text{NO}_3^-$ reduction rates | | <0.0001 | ns | ns | ns | ns |
| $\text{NO}_x$ reduction rates | | | ns | ns | ns | ns |
| $\text{NO}_2^-$ dynamics | | | | ns | ns | ns |
| $\text{N}_2\text{O}$ dynamics | | | | | ns | ns |
| Methanol consumption rates |  |  |  |  |  | ns |

ns: nonsignificant.
All correlations are directly proportional.
i-Methanol and i- $\text{NO}_3^-$ : Initial methanol and initial $\text{NO}_3^-$ concentrations, respectively.
Data obtained after Covid-19 (C6, C7 and C8, Period 2) were not included in the analysis.

### 3.3 Bacterial community

The evolution of the bacterial community of the *RR biofilm* during the different operating conditions were assessed by performing PCR amplification of the 16S rRNA genes and by analysing the composition of the amplicons by DGGE and amplicon sequencing. The PCR-DGGE profiles showed striking differences between the *Original Biofilm* sample and the samples taken from the reactor (Fig. 3A). The profile of the D0 sample revealed some similarities with the profiles derived from samples taken during the operating conditions. The PCR-DGGE profiles of these later samples did not reveal substantial changes between them, which suggesting that the bacterial community was stable during the operating conditions.

**Figure 3.**
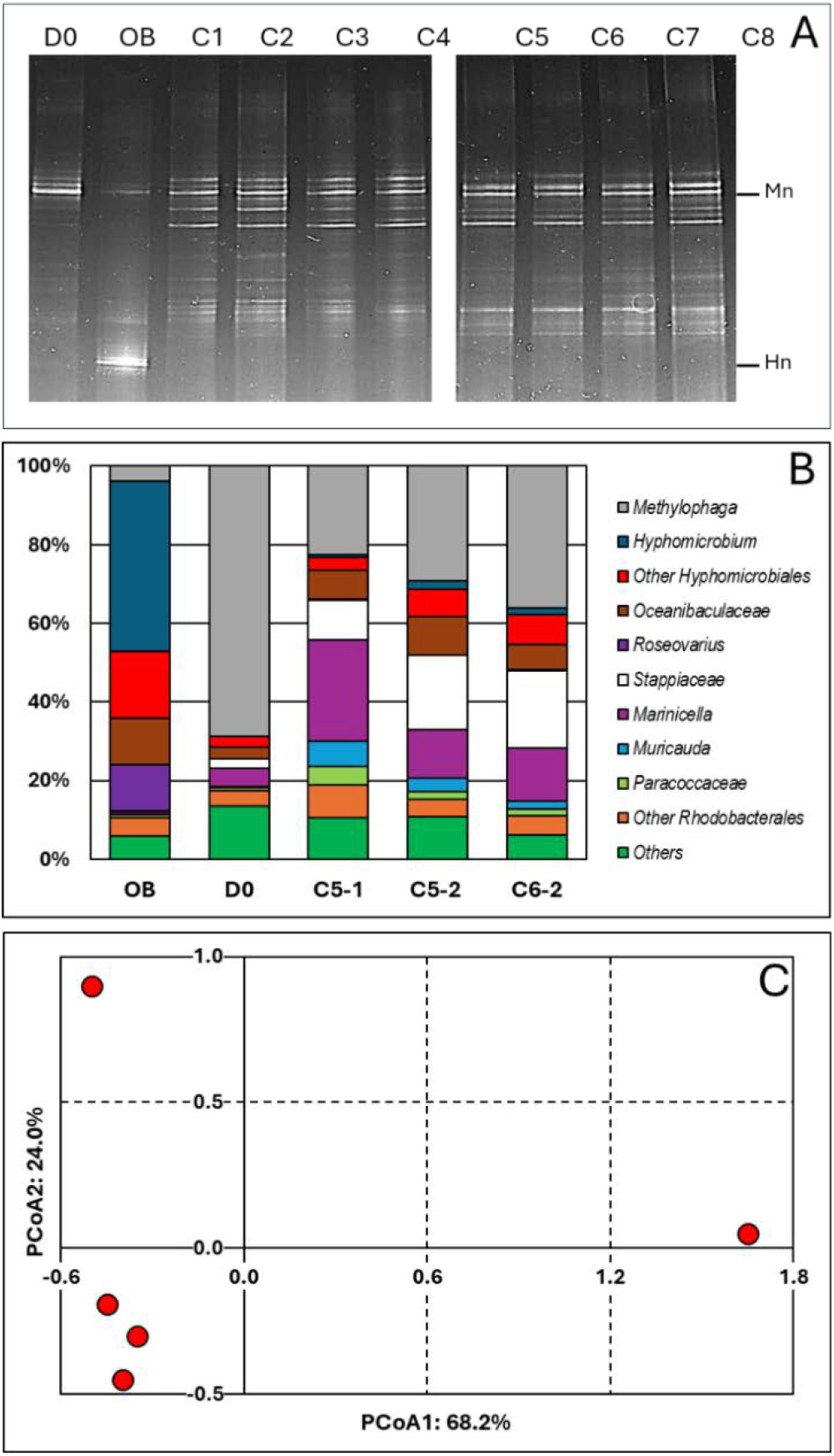
Bacterial profiles in the biofilm samples. <u>Panel A.</u> DNA was extracted from biofilm samples taken from the reactor at Day 0 (D0) and during Period 1, conditions 1 to 8 (C1 to C8), and taken from the Original Biofilm (OB). The V3 region of the 16S rRNA genes were amplified with one primer having a GC clip, and the resulting amplicons were resolved by denaturing gradient gel electrophoresis. Mn and Hn: Locations of the migration of the V3 sequences of *Methylophaga nitratireducenticrescens* and *Hyphomicrobium nitrativorans*, respectively. <u>Panel B.</u> DNA were extracted from the recirculating reactor at D0 and during the operating conditions: Condition 5 (P1 and P2; C5-1, C5-2) and Condition 6 (P2; C6-2). For comparison, we added data from the Original Biofilm (OB). Part of the 16S rRNA gene was amplified and the resulting amplicons were sequenced. Taxonomic affiliations of the resulting sequences were carried out, and their proportions derived. The most relative abundant taxa are illustrated. <u>Panel C</u>. Principal coordinate analysis (PCoA) of the bacterial profiles of the five biofilm samples was carried out using Bray-Curtis distance calculation.

We performed amplicon 16S sequencing on total DNA extracted from the *RR biofilm* samples. The first sample was taken at D0 (C/N of 3.0; 21.4 mM NO_3_^−^, 0.3% methanol). Based on the DGGE profiles, we have chosen three representative samples during the operating conditions: two were taken from the reactor operated under Condition 5 (21.4 mM NO_3_^−^ and 0.45% methanol: C/N of 4.5) during P1 (C5-1; Day 30) and P2 (C5-2; Day 120). The third representative sample was taken from the reactor operated under Condition 6 (21.4 mM NO_3_^−^ and 0.75% methanol; C/N of 7.5) during P2 (C6-2; Day 200) after the Covid break. For comparison, we added data that we obtained from the *Original Biofilm* of the Biodome denitrification system (estimated C/N of 0.86) (Labbé *et al*., 2003).

We identified forty-two taxa across the five biofilm samples and determined their proportions in the biofilm samples. Figure 3B illustrates the proportions of the most abundant taxa observed in the samples. Principal coordinate analysis (PCoA) showed that the bacterial profiles of the C5-1, C5-2 and C6-2 biofilm samples were similar; the profiles of the D0 and *Original Biofilm* samples differed from each other, and from the other ones (Fig. 3C). These results concur with the PCR-DGGE results. Because of the unicity of the sample (no replicate), we could not perform statistical analysis such as PERMANOVA to determine significance in variations between the five biodiversity profiles. However, it is obvious that the bacterial profile of the *Original Biofilm* is totally different than these of the four *RR biofilm* samples.

Thus, higher proportions of *Hyphomicrobiales, Roseovarius* spp. and *Oceanibaculaceae*, and much lower proportion of *Methylophaga* spp. were found in the *Original Biofilm* samples compared to the four *RR biofilm* samples. In the D0 biofilm sample, we noticed much higher relative abundance of *Methylophaga* spp. (69%), This result corresponds with the DGGE profile of the D0 biofilm sample where an intense DNA fragment associated to *M. nitratireducenticrescens* was observed. For the C5-1, C5-2 and C6-2 biofilm samples, increases in the relative abundance of *Marinicella* spp., *Stappiacaea* spp., *Muricauda* spp. and *Oceanibaculaceae* were observed compared to the D0 biofilm sample with approximately 50% decrease in relative abundance of *Methylophaga* spp. The C5-1, C5-2 and C6-2 biofilm samples had higher alpha diversity indexes compared to the *Original Biofilm* (Table 3). Because of high proportion of *Methylophaga* spp. in the D0 biofilm sample, the alpha diversity indexes were lower than those in the four other biofilm samples. Regarding the beta diversity (Table 3), examining the common taxa between samples, taxa found in the *RR biofilm* samples shared the greatest number of taxa. All these results showed that the original bacterial community evolved in diversification in the recirculating reactor from Day 0, and then was relatively stable during the operating conditions.

**Table 3:**
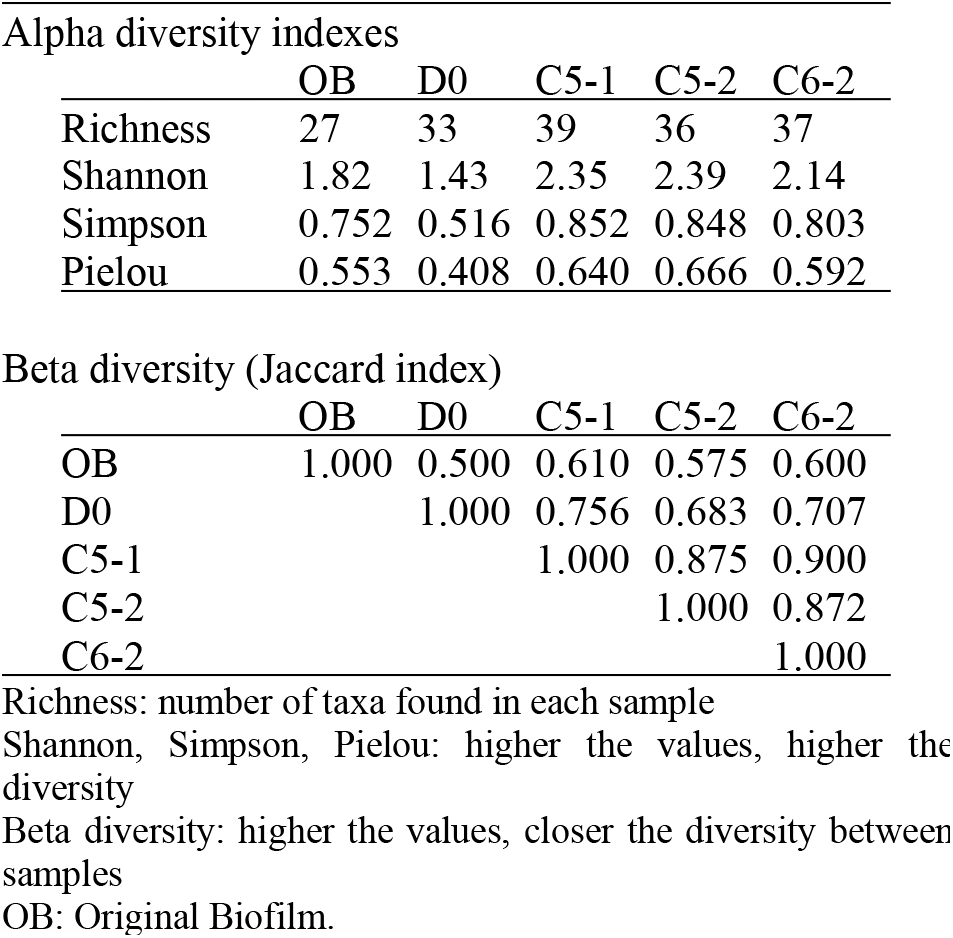
Diversity indexes of taxa deduced from 16S amplicon sequencing.

### 3.4 Metatranscriptome analysis

We derived metatranscriptomes of the microbial community in four *RR biofilm* samples to determine (1) the relative transcript profiles of *M. nitratireducenti-crescens* strain GP59, the main denitrifier in the *RR biofilm*; and (2) the potential metabolic pathways in function in the *RR biofilm*.

The metatranscriptomes were derived during P1 from: Condition 1 (C1-1; C/N of 1.5; 10.7 mM NO_3_^−^, 0.075% methanol; Day 4), Condition 4 (C4-1; C/N of 3.0; 21.4 mM NO_3_^−^, 0.3% methanol; Day 24). Condition 6 (C6-1; C/N of 7.5; 21.4 mM NO_3_^−^, 0.75% methanol; Day 36), and Condition 8 (C8-1; C/N of 3; 42.8 mM NO_3_^−^, 0.6% methanol; Day 89). Condition 1 and Condition 6 were chosen as the lowest (1.5) and highest (7.5) C/N applied to the reactor, respectively. Although Condition 4 and Condition 8 had the same C/N (3.0), they were adjusted with distinct levels of NO_3_^−^ and methanol. In addition, we derived the metatranscriptome of the *Original Biofilm* for comparison.

#### 3.4.1 Relative transcript profiles of *M. nitratiredu-centicrescens* strain GP59

The proportion of transcript reads associated to strain GP59 in the metatranscriptomes were similar in the four *RR biofilm* RNA samples (average 7.15%; Table 4), which is four times higher than that in the *Original Biofilm* RNA sample. Because of the unicity of the sample (no replicate), we could not perform statistical analysis such as PERMANOVA to determine significance in the variations between the four expression profiles.

**Table 4.**
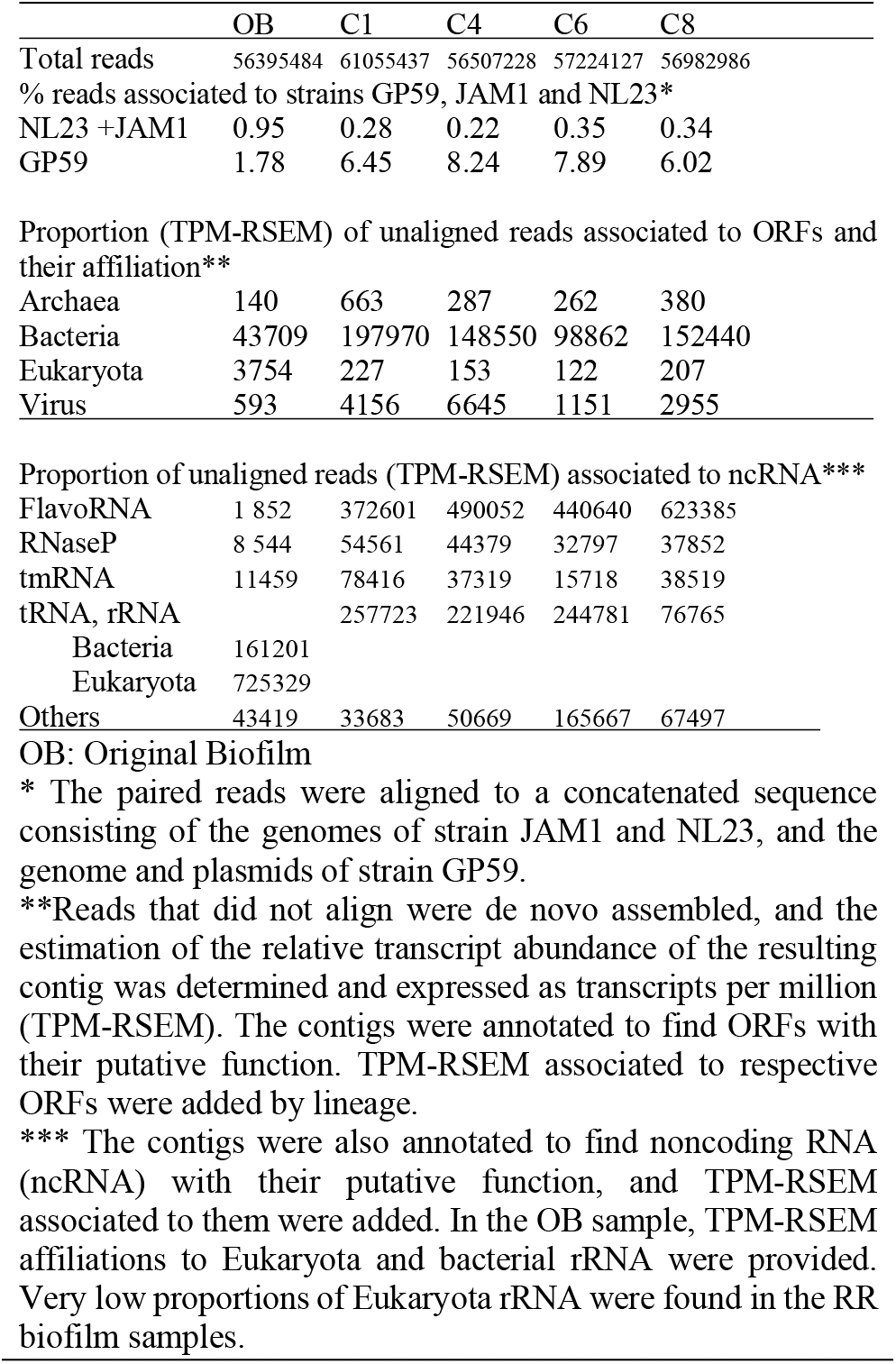
Proportion of transcriptomic reads affiliated to lineage or to ncRNA.

We took this opportunity to compare the relative gene expression profiles of strain GP59 derived from the four *RR biofilm* RNA samples and from the *Original Biofilm* RNA sample with profiles derived from our previous studies. These include:

1. The *Original Biofilm* cultured in vials under batch-static conditions in artificial seawater (ASW, 2.75% NaCl), with two different NO_3_^−^ concentrations and temperatures: «300N23C» (21.4 mM NO_3_^−^, 0.15% methanol, C/N of 1.5, 23°C); and «900N30C» (64.2 mM NO_3_^−^, 0.45% methanol, C/N of 1.5, 30°C) (Payette *et al*., 2019, Villemur *et al*., 2019).
2. The *Original Biofilm* cultured in vials under batch-static conditions in low salt concentration ASW: «0%NaCl» (ASW with 0% NaCl, 21.4 mM NO_3_^−^, 0.15% methanol, C/N of 1.5, 23°C) (Payette *et al*., 2019, Villemur *et al*., 2019).
3. Planktonic cultures (triplicate) of strain GP59 cultured in a *Methylophaga*-specific culture medium (*Methylophaga*-1403, 0.3% methanol; C/N of 3) under oxic conditions with NO_3_^−^ (21.4 mM, PlkON) or without NO_3_^−^ (PlkO), and under anoxic conditions with 21.4 mM NO_3_^−^ (PlkAN) (Lestin & Villemur, 2024).

All these cultures differ in the culture media (IO, ASW, low salt ASW, *Methylophaga*-1403), C/N, the operating mode (batch-static, continuous, recirculating sequential-batch), and the physiology (biofilm, planktonic). The goal of this analysis was to reveal metabolisms in strain GP59 that could have been impacted by the distinct culture conditions. Principal coordinate analysis (PCoA) clearly showed distinct grouping of each category of cultures, explained by 68.5% variations (Fig. 4). PERMANOVA were performed with the planktonic cultures (triplicate: *Methylophaga*-1403 medium, anoxic or oxic, presence or absence of NO_3_^−^, planktonic) and the four *RR biofilm* samples by assuming that the relative transcript profiles of the *RR biofilm* RNA samples were similar enough to be considered as replicates (IO medium, anoxic, biofilm, presence of NO_3_^−^). The results showed, as perceived in Figure 4, that the medium and the physiology (IO [biofilm] vs M1403 [planktonic]; r^2^: 0.652, p: 0.001), and the operating conditions (oxic vs anoxic; r^2^: 0.144, p: 0.001) had substantial effect on the relative transcript profiles of strain GP59.

**Figure 4.**
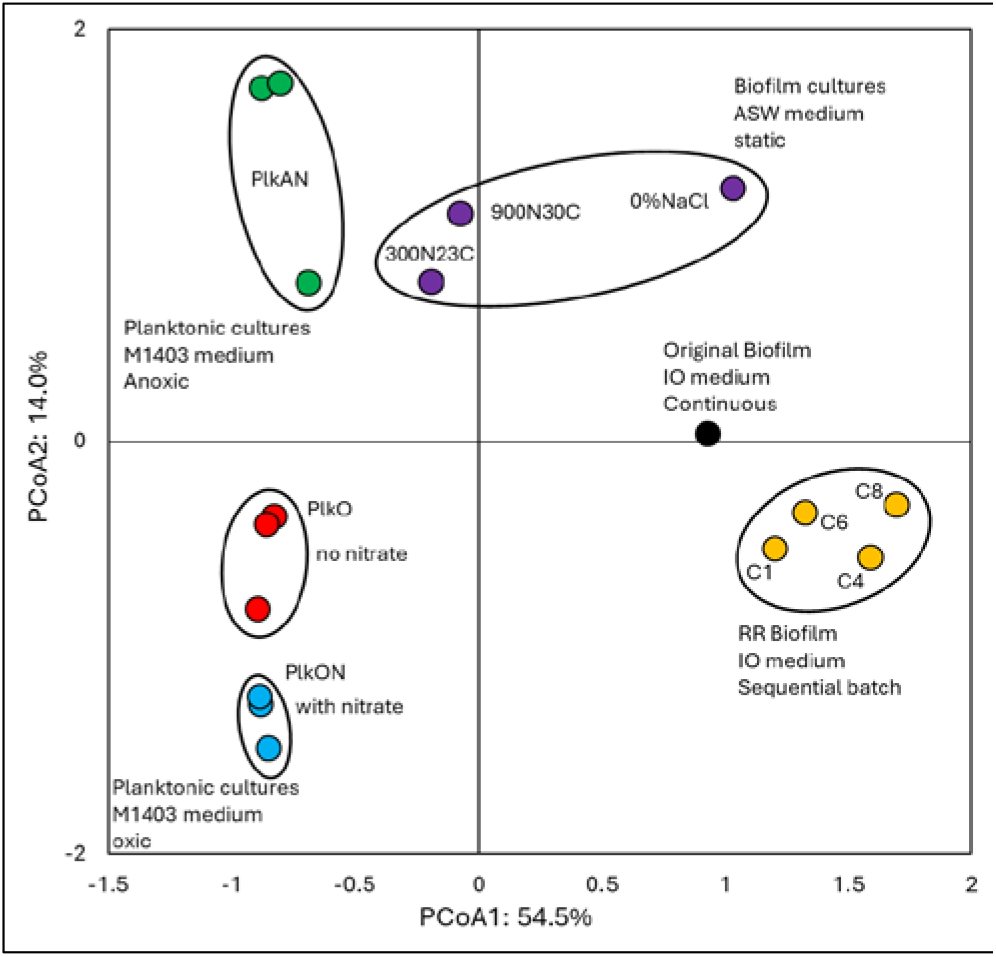
Principal coordinate analysis of the relative transcript profiles of strain GP59 in different cultures. Metatranscriptomes of the four RR biofilm samples, and the Original Biofilm were performed, and the relative transcript profiles of the 3204 genes of strain GP59 were derived. The strain GP59 gene expression profiles from our previous studies were also included in PCoA. These profiles were (1) the Original Biofilm cultured in ASW medium under different conditions (300N23C; 900N30C, 0%NaCl), and pure cultures of strain GP59 cultured in Methylophaga 1403 medium under oxic or anoxic conditions (PlkO, PlkON, PlkAN). See the text for complete description. PCoA by percentage difference (Bray-Curtis distance) were performed with the Canoco software for ordination version 5.15.

We looked at the relative transcript levels of genes involved in the N cycle. Thirty-three genes or gene clusters were selected for their involvement in denitrification, N assimilation, regulation and NO-responses. PCoA was performed to derive the preferential expression pattern of these genes in the different types of cultures. Figure 5 illustrates the PCoA pattern that explained 78% variations. These genes and cultures segregated in three specific quadrants: (i) bottom left quadrant with the planktonic oxic cultures (PlkO, PlkON); (ii) bottom right quadrant with the 0%NaCl biofilm (non-marine conditions) and the *Original Biofilm*; and (iii) top quadrants with the planktonic anoxic cultures (PlkAN) and the *RR biofilm* samples

**Figure 5.**
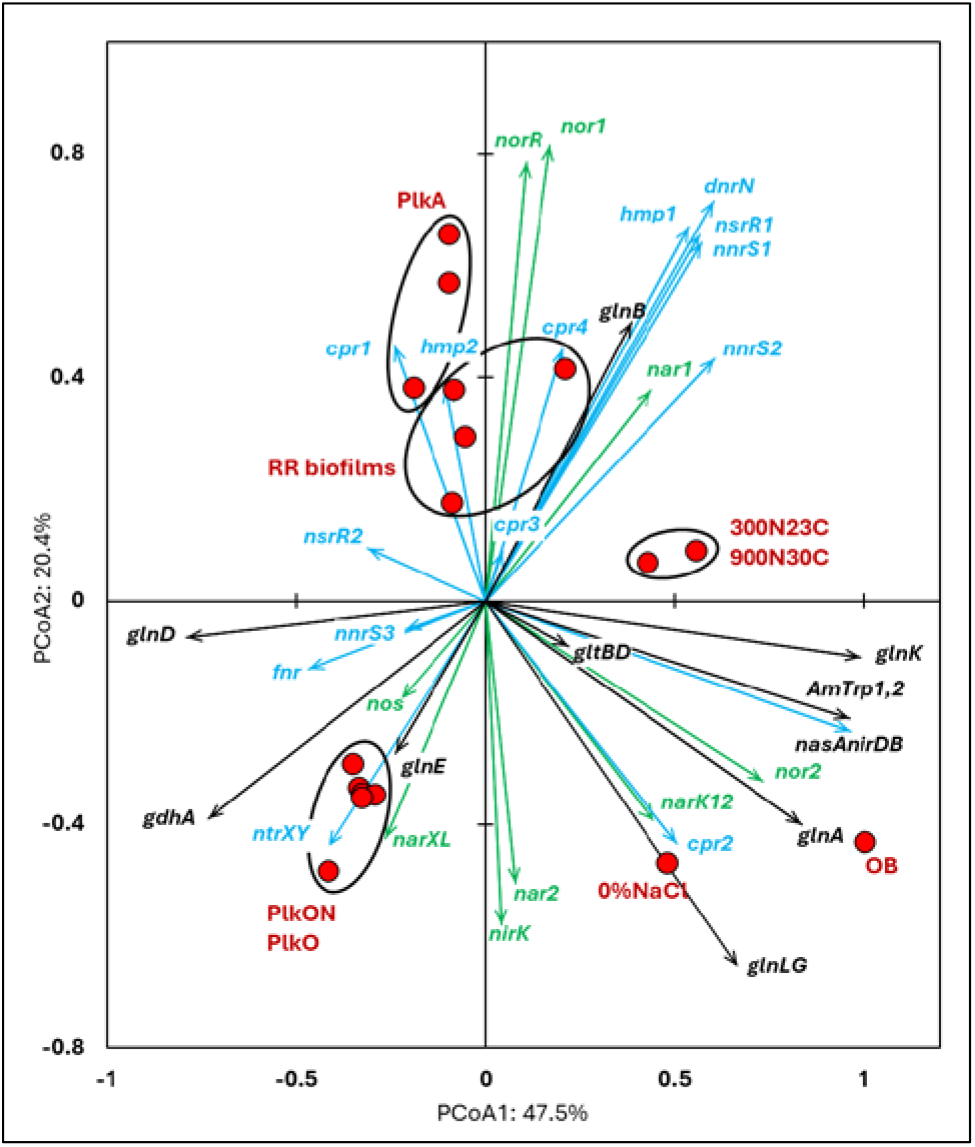
Principal coordinate analysis of the relative transcript profiles of genes involved in the N cycle of strain GP59 in different cultures. Thirty-three genes or gene clusters were identified in strain GP59 genome to be involved in denitrification, N assimilation, regulation and NO-responses. Their relative transcript levels derived from the biofilm metatranscriptomes, or from pure culture transcriptomes (see Figure 4 legend) was used in PCoA. PCoA by percentage difference (Bray-Curtis distance) were performed with the Canoco software for ordination version 5.15. *narXL*: NO_3_^−^/NO_2_^−^ transcription regulator; *nar1*: NO_3_^−^ reductase 1 operon with two *narK* transporters; *narK12f*: NO_3_^−^ transporter; *nar2*: NO_3_^−^ reductase 2 operon; *nirK*: NO-forming NO_2_^−^ reductase; *nor1*: nitric oxide reductase 1 gene cluster; *norRE*: nitric oxide reductase transcription regulator; *nor2*: nitric oxide reductase 2 gene cluster; *nos*: nitrous oxide reductase gene cluster; *nasAnirDB*: assimilatory NO_3_^−^/NO_2_^−^ reductases; cAMP CRP1,2,3,4: represent four genes encoding Crp/Fnr family transcriptional regulators; *fnr*: fumarate/NO_3_^−^ reduction transcriptional regulator Fnr; *nnrS1*,*2*,*3*: represent three genes encoding protein involved in response to NO; *nsrR1*,*2*: represent two genes encoding nitric oxide-sensitive transcriptional repressors; *dnrN/ytfE*: iron-sulfur cluster repair protein; *hmp1*,*2*: represent two genes encoding Nitric oxide dioxygenases; *ntrXY*: Nitrogen regulation protein; AmTrp1,2: represent two genes encoding Ammonium transporters; *gltBD*: glutamate synthase; *gdhA*: glutamate dehydrogenase; *glnE*: [glutamate--ammonia-ligase] adenylyltransferase; *glnB*: P-II family nitrogen regulator; *glnK*: P-II family nitrogen regulator; *glnD*: [protein-PII] uridylyltransferase; *glnA*: glutamate--ammonia ligase.

Several genes in the N assimilation pathways (ammonium transporters, *nasAnirDB, glnA, glnK, glnLG*) covaried in their relative expression with the *Original Biofilm* and the 0%NaCl biofilm (Fig. 5). In addition, the denitrification genes *nor2* and *narK12f* covaried with these biofilm cultures.

The regulatory genes, *fnr, ntrXY* and *narXL*, and the denitrification genes *nos, nirK* and *nor2* covaried in relative transcripts levels with the PlkO and PlkON (oxic) cultures in opposition with *nar1, nor1* and its associated regulatory genes *norRE*, which covaried with the PlkAN cultures and the *RR biofilm* (Fig. 5). Furthermore, the relative transcript levels of *nnrS1, nsrR1* and *dnrN*, which are adjacent genes in strain GP59 genome, and *hmp1* and *nnrS2* covaried with the PlkAN cultures, the *RR biofilm* and the two ASW biofilm cultures 300N23C and 900N30C. These genes encode putative regulators or proteins with NO-response function and may be related to higher NO production in these cultures.

We looked then to the relative expression profiles of genes involved in the C cycle. Among the 112 identified genes or gene clusters, we have chosen 17 genes or gene clusters for PCoA, in which their relative transcript levels varied more than five times (significative differences as deduced by EdgeR analysis with FDR <0.05) between at least two conditions. We also included in the PCoA the relative transcript profiles of the eleven riboswitches (Ribo1 to Ribo11) identified in strain GP59 genome.

Ribo1 to 7 are c-di-GMP riboswitches; Ribo9 and 10, cobalamin riboswitches adjacent to each other and flanked by genes involved in cobalamin synthesis; Ribo8 and Ribo11, thiamine pyrophosphate and S-adenosyl methionine riboswitches, respectively.

We found that covariation occurred between the relative expression of the four *xoxF* genes (encoding putative lanthanide [Ln^3+^]-dependent methanol dehydrogenases) and the four *RR biofilm* samples (Fig. 6). Also, two genes involved in CO2 assimilation, encoding the pyruvate carboxylase (*pycAB*; HCO_3_^−^ to oxaloacetate) and the carbonic anhydrases (*cynT*; dissolved CO_2_ to HCO_3_^−^), co-varied with the *RR biofilm* samples. The relative expression profiles of the methanol dehydrogenase gene cluster (*mxa*) and one of the formate dehydrogenase gene clusters (*fdh2*) covaried with PlkO, PlkON cultures. Interestingly, the relative expression profiles of 10 out of 11 riboswitches covaried with the *RR biofilm* samples (Fig. 6). The other riboswitch (Ribo5) covaried with the PlkAN cultures.

**Figure 6.**
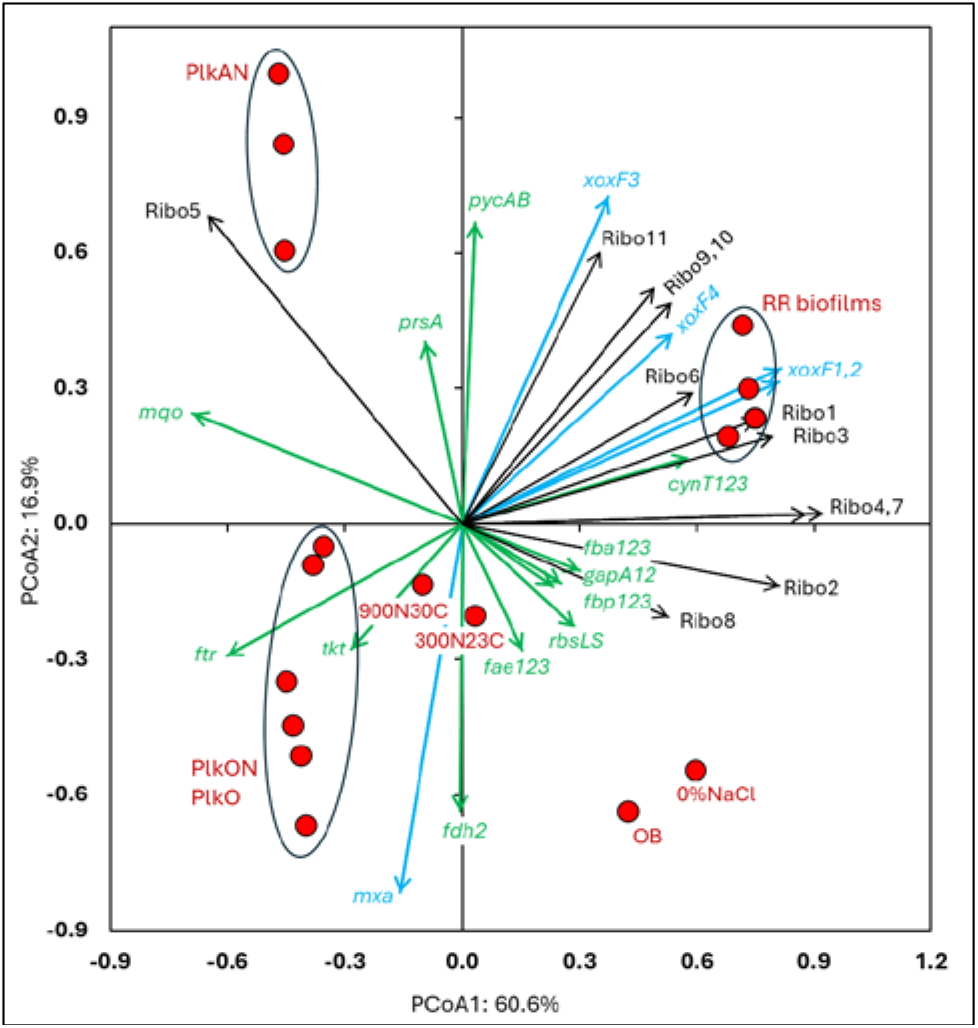
PCoA of the relative transcript profiles of genes involved in the Carbon cycle and riboswitches of strain GP59 in different cultures. One hundred twelve genes or gene clusters were identified in strain GP59 genome in the C cycle. Among them, seventeen in which their relative transcript levels varied more than five times between at least two conditions were chosen for PCoA. The relative transcript profiles of the eleven riboswitches (Ribo1 to Ribo11) were also included in PCoA. PCoA by percentage difference (Bray-Curtis distance) were performed with the Canoco software for ordination version 5.15. *mxa*: methanol dehydrogenase; *xoxF1*,*2*,*3*,*4*: represent four genes encoding putative lanthanide [Ln3+]-dependent methanol dehydrogenases; *fae1*,*2*,*3*: represent three genes encoding 5,6,7,8-tetrahydromethanopterin hydro-lyases; *ftr*: formylmethanofu-ran--tetrahydromethanopterin N-formyltransferase; *fdh2*: formate dehydrogenase; *tkt*: transketolase; *prsA*: ribose-phosphate pyrophosphokinase; *rbcLS*: ribulose-bisphosphate carboxylase; *fbp1*,*2*,*3*: represent three genes encoding fructose-1,6-bisphosphatases; *fbaA1*,*2*,*3*: represent three genes encoding fructose-bisphosphate aldolases; *gapA1*,*2*: represent two genes encoding glyceraldehyde 3-phosphate dehydrogenases; *pycAB*: pyruvate carboxylase; *mqo*: malate dehydrogenase (quinone); *cynT1*,*2*,*3*: represent three genes encoding carbonic anhydrases; Ribo1 to Ribo7: c-di-GMP riboswitches, Ribo8: thiamine pyrophosphate riboswitch, Ribo9 and 10: cobalamin riboswitches, Ribo11: S-adenosyl methionine riboswitch.

#### 3.4.2 Diversity of transcriptionally active bacteria

We analysed reads that were not associated to *M. nitratireducenticrescens* and *Hyphomicrobium nitrativorans* to determine taxa that were the most transcriptionally active in the five biofilm samples, and to assess the potential metabolic pathways used by these taxa. Sequencing reads from the metatranscriptomes were aligned to the genomes and plasmids of *M. nitratireducenticrescens*, and of *Hyphomicrobium nitrativorans*. Between 2.7% to 8.5% reads aligned to these genomes in the five biofilm RNA samples (Table 4). Reads that did not align were *de novo* assembled, and the relative transcript levels of the resulting contigs (expressed as TPM-RSEM) was determined by aligning reads to them. The contigs were analyzed for the presence of ORF and processed to databases to determine their putative function and their most probable taxonomic affiliation.

We identified nine-three taxa across the 5 biofilm samples, among which sixty-three were found in all samples. The proportion of transcripts affiliated to *Methylophaga* spp. (excluding *nitratireducenticrescens*) accounted for around 49% in the *Original Biofilm* RNA sample but decreased substantially (0.19 to 1.3%) in the *RR biofilm* RNA samples (Table 5). Seventeen taxa low in proportion in the *Original Biofilm* RNA sample showed substantial increases (one to four orders of magnitude) in their relative transcript levels in at least one of the *RR biofilm* RNA samples, whereas those affiliated to the genera *Sedimenticola, Dechloromarinus, Cycloclaticus* and *Leptolyngbya* presented one to three orders of magnitude decreases in all the *RR biofilm* RNA samples (Table 5). Relative transcript levels affiliated to the genera *Bradymonas, Roseovarius* and Cand. *Promineofilum* remained at the same proportion (within one order of magnitude) in all biofilm RNA samples (Table 5).

**Table 5.**
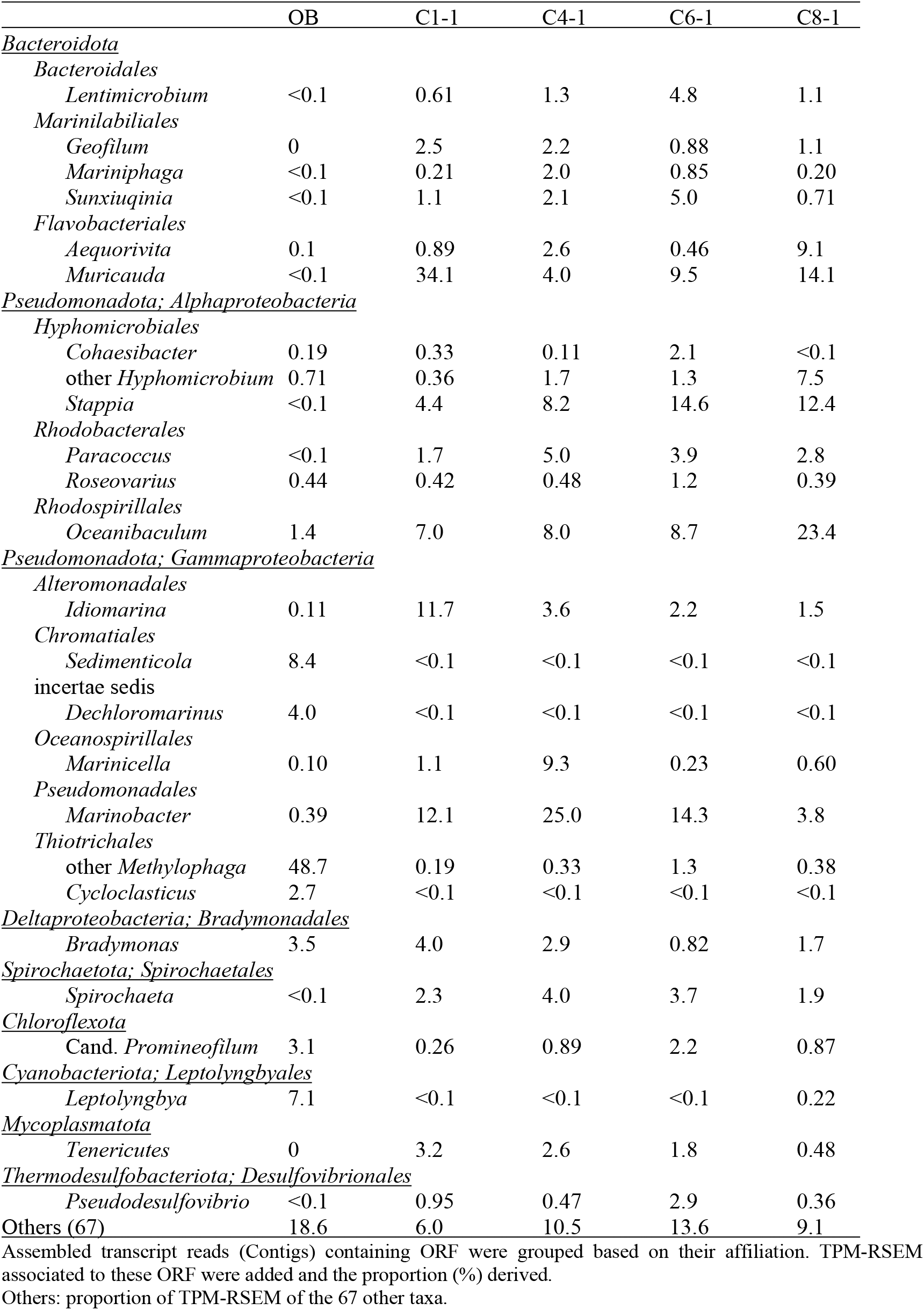
Taxonomic affiliation of contigs with ORF and their proportions.

High proportions of contigs associated with ncRNA were found in all samples (Table 4). These ncRNA were related to rRNA, RNAseP (RNA component; RPR) and tmRNA. We also found contigs in the four RR biofilm RNA samples that mapped to an intergenic region found in several Flavobacteriale genomes (here named FlavoRNA). The analysis of these sequences will be subjected to another article in BioRxiv.

#### 3.4.3 Functional diversity

Contigs with ORF that were grouped based on their taxonomic affiliation provided an estimate of the metabolic activities occurring in the associated taxa in the biofilm. Among the ninety-three identified taxa, twenty-five taxa, in which their relative transcript levels of grouped contigs were >2% TPM-RSEM in at least one RNA sample, were examined for putative metabolic pathways in action in these taxa (Table 5). Genes involved in denitrification (the four reductases) and in the carbon cycle (Fig. 7) were more closely examined as these pathways could explain the potential of these taxa to harvest energy and to assimilate the C1 carbon for their growth and maintenance. The chosen bacterial pathways were the following.

**Figure 7.**
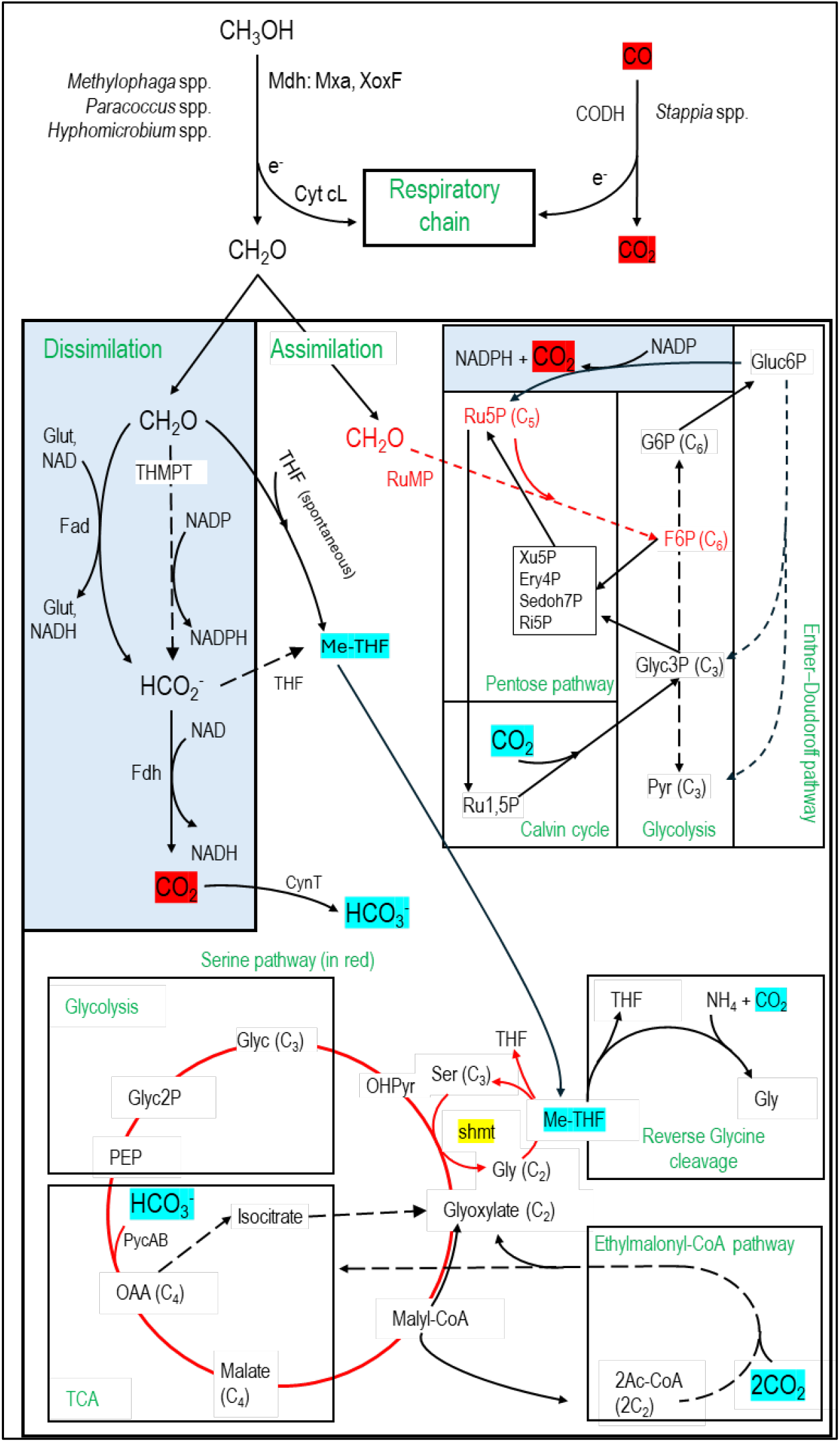
Carbon assimilation and dissimilation pathways of methanol and formaldehyde. Pathways were retrieved from KEGG and MetaCyc databases. Dashed arrows refer to pathways with multiple steps. C2, C3, C4, etc. refer to the number of carbons of the corresponding molecules. MeTHF: Methylene tetrahydrofolate. Ery4P: Erythrose-4P. Ru5P: Ribulose-5P. Ru1,5P: Ribulose-1,5P. Xu5P: Xylulose-5P. Ri5P: Ribose-5P. Sedoh7P: Sedoheptulose-7P. G6P: Glucose-6P. F6P: Fructose-6P. Pyr: Pyruvate. Glyc: Glycerate. Gluc6P: Gluconate-6P. OAA: Oxaloacetate. OHPyr: Hydroxypyruvate. Ac-CoA: Acetyl-CoA. shmt: Serine hydroxymethyltransferase. PEP: Phosphoenolpyruvate carboxylase. THMPT: Tetrahydromethanopterin. Glut: Glutathione. Mxa: Methanol dehydrogenase. XoxF: lanthanide [Ln3+]-dependent methanol dehydrogenase. CODH: carbon monoxide dehydrogenase. Cyt: Cytochrome. PycAB: pyruvate carboxylase. CynT: carbonic anhydrase. Dissimilatory pathways are blue shaded.

1. Methanol oxidation to formaldehyde by the methanol dehydrogenase (Mdh), which provides gain of reductive equivalents by reducing cytochromes for the respiratory chain (Anthony, 1982).
2. Formaldehyde dissimilation pathways to formate by formaldehyde dehydrogenases (Fad; glutathione-dependent or independent) or by the tetrahydro-methanopterin (THMPT) pathway, and formate to CO_2_ by formate dehydrogenases (Fdh). Gain of reductive equivalents (NADH) was obtained in these dissimilation pathways (Anthony, 1982, Vorholt, 2002).
3. Formaldehyde assimilation pathways through the ribulose monophosphate (RuMP). Associated with RuMP is the Entner–Doudoroff pathway, with carbon assimilation to pyruvate and glyceraldehyde3P, or carbon dissimilation by regenerating Ru5P with a lost of CO_2_ but gain of energy (NADPH) (Anthony, 1982, Vorholt, 2002).
4. Formaldehyde assimilation pathway through the methyl transfer (spontaneous) to tetrahydrofolate (THF) generating methylene-THF (meTHF), which is in turn incorporated in the biomass by the serine pathway or putatively by the reverse glycine cleavage system (RevGCS) (Bar-Even *et al*., 2013). Formate can be transformed to meTHF by the THF pathway (Anthony, 1982).
5. The Calvin cycle and the ethylmalonyl-CoA pathway (Alber, 2011) were also examined as associated with the pentose pathway or the Serine pathway. These pathways can provide additional carbon in the biomass through CO_2_ incorporation (Anthony, 1982).
6. Nineteen taxa contain transcripts associated with denitrification genes (Table 6), among which genes encoding the four reductases were found in the genera *Hyphomicrobium, Marinobacter, Methylophaga, Sedimenticola* and *Stappia*. In addition to *Methylophaga* spp. and *Hyphomicrobium* spp., *Paracoccus* spp. was the only other taxon with transcripts associated to methanol dehydrogenase (Table 6). The formaldehyde dissimilation pathways (Fad, THMPT, Fdh; Fig. 7) were found in eighteen taxa, among which three have THMPT and Fdh, and eight have Fad and Fdh (none have the three; Table 6). Only *Methylophaga* spp. use the RuMP pathway (Fig. 7). The THF pathway and the serine pathway (Fig. 7) were found in eleven and ten taxa, respectively (six taxa contain both; Table 6), whereas GCS was found in all taxa. The Calvin cycle, and ethylmalonyl-CoA pathway (Fig. 7), where found in six and four taxa, respectively (Table 6).

**Table 6:** Transcripts associated with genes involved in denitrification, and in formaldehyde assimilation and dissimilation pathways.

| Taxon | Denitrification | Mdh | Dissimilation |  |  | Assimilation |  |  |  |  |  |
| --- | --- | --- | --- | --- | --- | --- | --- | --- | --- | --- | --- |
|  |  |  | Fad | THMPT | Fdh | THF | RuMP | Calvin | Ser | EM | ED |
| *Aequorivita* * | *nir, nor, nos* | - | - | - | - | + | - | - | - | - | - |
| *Bradymonas* * | *nap, nir* | - | + | - | + | - | - | - | + | - | - |
| *Cand. Promineofilum* | *nir, nos* | - | - | - | + | - | - | - | - | - | - |
| *Cohaesibacter* | *nir, nor, nos* | - | + | - | + | - | - | - | - | - | - |
| *Cycloclasticus* * |  | - | + | - | + | - | - | - | + | - | - |
| *Dechloromarinus* * | *nar, nor, nos* | - | + | - | + | - | - | + | + | - | - |
| *Geofilum* | *nor* | - | - | - | - | + | - | - | - | - | - |
| *Hyphomicrobium* | *nap, nir, nor, nos* | + | - | + | + | + | - | - | + | + | - |
| *Idiomarina* * | *nar, nir, nor* | - | + | - | - | - | - | - | - | - | - |
| *Lentimicrobium* |  | - | - | - | + | - | - | - | - | - | - |
| *Leptolyngbya* * |  | - | + | - | - | - | - | + | - | - | - |
| *Marinicella* | *nir, nor* | - | + | - | - | - | - | - | - | - | - |
| *Mariniphaga* |  | - | - | - | - | - | - | - | - | - | - |
| *Marinobacter* * | *nar, nir, nor, nos* | - | + | - | + | - | - | - | + | - | + |
| *Methylophaga* | *nar, nir, nor, nos* | + | - | + | + | + | + | + | - | - | + |
| *Muricauda* | *nor, nos* | - | - | - | - | + | - | - | - | - | - |
| *Oceanibaculum* | *nar* | - | + | - | + | + | - | - | + | - | - |
| *Paracoccus* | *nar, nir, nor, nos* | + | + | - | + | + | - | + | + | + | + |
| *Pseudodesulfovibrio* * | *nar* | - | - | - | + | - | - | + | - | - | - |
| *Roseovarius* * | *nir, nor, nos* | - | + | - | + | + | - | - | + | + | - |
| *Sedimenticola* * | *nar, nir, nor, nos* | - | - | + | + | + | - | + | + | - | - |
| *Spirochaeta* * |  | - | - | - | - | - | - | - | - | - | - |
| *Stappia* * | *nap, nir, nor, nos* | - | + |  | + | + | - | - | + | + | + |
| *Sunxiuqinia* | *nir, nor* | - | - | - | - | + | - | - | - | - | - |
| *Tenericutes* |  | - | - | - | - | - | - | - | - | - | - |

\*Pathways were identified by the KEGG mapper reconstruct analysis with data from the metatranscriptomes, and by the KEGG pathways analysis with a strain representative of the taxon when available. +: presence; -: absence.
See Figure 7 for pathway description.
-Mdh: Methanol dehydrogenase (*mxh* mainly)
-Fad: Formaldehyde to formate by the formaldehyde dehydrogenase (either glutathione-dependent or -independent)
-THMPT: Process of formaldehyde to formate through the Tetrahydromethanopterin (THMPT) pathway
-Fdh: Formate dehydrogenase
-THF: Process of formate to Methylene tetrahydrofolate (meTHF) through the THF pathway
-RuMP: Formaldehyde assimilation by the Ribulose-5P pathway
-Calvin: Calvin cycle here is defined as taxon containing transcripts encoding the Prk and Rubisco enzymes
-Ser: Formaldehyde assimilation by the serine pathway

## 4 Discussion

### 4.1 C/N and the denitrification performance

Our results showed that different C/N have not significantly impacted the denitrification performance of the reactor (NOx/NO_3_^−^ reduction rates, NO_2_^−^ and N_2_ O dynamics), but did have an impact on the methanol consumption rates and the CO_2_ production rates. Such impact can be linked to the i-Methanol applied to the reactor, as these concentrations also correlated with the methanol consumption rates and the CO_2_ production rates. Interestingly, the i-NO_3_^−^ did positively correlates with the denitrification performance. Such correlation was not observed with the i-Methanol. Our results suggest that opposite effects on the denitrification performance occur between the i-NO_3_^−^ and the i-Methanol applied to the reactor, which translated into higher impact on the carbon pathways.

The reactor was able to reduce most, if not all, all i-NO_3_^−^ applied to the reactor after four operating days, although the highest concentrations (32.1 and 42.8 mM) generated higher levels of residual NO_2_^−^, which would be eventually reduced over time. The reactor did not generate ammonium, and negligible amount of N_2_ O was found at the end of the reaction phase. However, high proportion of residual methanol (47 to 91%) were found at the end of the denitrification process, which is not a desire outcome for a denitrification system. Releasing methanol in the effluent of a water treatment can be harmful for the ecosystem. For instances, Pungrasmi et al. (2013) showed that applying a C/N of 4 with methanol in their denitrification system gave the highest denitrification performance. However, because of the concentration of residual methanol exceeded toxic level for fish development, a C/N of 3.3 had to be applied in their system when treating the effluent of a recirculation (tilapia) aquaculture system (RAS).

Numerous studies reported a wide range of optimal C/N with diverse types of denitrification processes (Fu *et al*., 2022, Brozincevic *et al*., 2024). We noticed that the operating conditions, continuous versus batch, of the studied denitrification processes have an impact in the determination of the optimal C/N to get the best denitrification performance. Whereas batch operating conditions can be adjusted to get optimal C/N for complete denitrification and minimal addition of carbon source, the continuous operating conditions have to also deal with the hydraulic retention time (HRT) of the reactor. For instances, in a denitrification reactor treating a marine RAS (sea bass) operating under continuous conditions, Turno et al. (2018), showed that optimal C/N of 2.1 to 2.3 achieved 75% total nitrogen removal with minimal NO_2_^−^ and ammonia in the effluent. However, they found that adjusting the HRT for better denitrification performance decreased the denitrification rates. In our previous works reported by Labelle et al. (2005), we developed 100-L moving bed biofilm reactor (pilot plant) in the purpose of improving the configuration of the Biodome denitrification system. This reactor was connected to the effluent of the Biodome seawater aquarium tank where a similar denitrifying microbial community colonized the supports. Operating under continuous regime, the reactor reached 88% denitrification efficiency (NOx removal) at a C/N ratio of 2.9. Below this level, incomplete denitrification occurred. Also, cautions have to be taken to not reaching a level of NO_3_^−^ too low (< 10 mg-N NO_3_^−^/L) as sulfate reduction can occur under marine conditions.

### 4.2 Evolution of the bacterial community

The bacterial community established in the reactor evolved during the 31-week operating time has diverged substantially from *Original Biofilm* taken from the Biodome denitrification system in the proportion of the common taxa. These changes can be explained by the operating conditions: laboratory-scale sequential batch-mode recirculating reactor versus full-scale continuous operating mode in the Biodome denitrification system. Although both processes used the same medium (IO), the input water in the Biodome system had additional matter from animal and plant waste, or non-ingested food (Parent *et al*., 2002), and additional microorganisms from the water ecosystem (e.g. protozoa) (Laurin *et al*., 2008) that could have influenced its microbial community.

The results obtained with 16S amplicon sequencing revealed that high proportion of *Methylophaga* spp. established in the recirculating reactor during the acclimation phase (D0), and that afterwards the biofilm populations increased in diversity at the expense of *Methylophaga* spp. (although still composed high proportion of the bacterial community), in favor of heterotrophic bacteria such as the genera *Stappia, Marinobacter, Marinicella, Oceanibaculum* and *Paracoccus. Methylophaga* spp., *Hyphomicrobium* spp., *Paracoccus* spp., *Marinobacter* spp. and *Marinicella* spp. were found in other marine denitrification systems (Güven, 2009, Lu *et al*., 2014, Furukawa *et al*., 2016, Wang *et al*., 2023). Some members of these taxa have denitrification capacity such as *Stappia* spp., *Paracoccus* spp. and *Marinobacter* spp. Contrary to what was claimed by Duc et al. (2018) (and carried by other reports) about Shoji et al. (2014), the sulfur oxidizing bacteria *Marinicella* were not shown to carry NO_3_^−^ reduction to N_2_ by this report. Results from metatranscriptomes revealed however that *nirK* and *nor* genes were found affiliated to *Marinicella* spp. None of the *Marinicella* sp. sequenced genomes in GenBank contain the full set of denitrification reductases. *Stappia* spp. are chemoorganotrophic marine bacteria that can oxidize CO, and some species contain a gene for the large subunit of ribulose-1,5-bisphosphate carboxylase/oxygenase (RuBisCO) (*cbbL*) that may suggest the capability to couple CO utilization to CO_2_ fixation (Weber & King, 2007).

Potential metabolic pathways to carry formaldehyde transformation for carbon assimilation or dissimilation (Anthony, 1982, Vorholt, 2002, Yurimoto *et al*., 2005) may explain persistence and growth of heterotrophs in the reactor, as these pathways were found in the observed taxa. For instance, NADH are generated from the oxidation of formaldehyde and formate by respective dehydrogenases. Formaldehyde is produced by the action of the methanol dehydrogenase, carried by *Methylophaga* spp., *Paracoccus* spp. and *Hyphomicrobium* spp., at the periplasm (Anthony, 2004). Because of the high proportion of *Methylophaga* spp. and high level of methanol, it is possible that excess of formaldehyde diffused in the biofilm and was available to the other organisms because of their promiscuity (Chistoserdova & Kalyuzhnaya, 2018). The emergence of these heterotrophic bacteria could also be the result of subsequent colonization of the biofilm after the early colonization of the supports by the methylotrophs such as *Methylophaga* spp., providing complex carbon to the heterotrophs such as carbohydrates in the extracellular polymeric substance of the biofilm matrix or organic waste from dead cells as the biofilm was getting older.

### 4.3 Relative expression profile of strain GP59

High proportions (22.5 to 68.9%) of 16S rRNA gene sequences affiliated to *Methylophaga* spp. were found in the *RR biofilm* samples, 6 to 18 times higher than in the *Original Biofilm* (3.7%). However, despite this low level in the *Original Biofilm*, transcript reads associated to strain GP59 and other *Methylophaga* species in the *Original Biofilm* represented together around 50% of the overall relative transcript profile of the *Original Biofilm*, which suggests that *Methylophaga* spp. were more active in the *Original Biofilm* than the other taxa. For instance, transcript reads associated to *Hyphomicrobium* spp. in the *Original Biofilm* RNA sample represented < 1% relative transcript levels, where it represented > 40% in 16S rRNA gene sequences in the DNA sample. Although relative transcript levels of strain GP59 had 4-fold increases in average in the *RR biofilm* RNA samples, these levels associated to the other *Methylophaga* species were almost negligible, suggesting that strain GP59 has better adapted to the recirculating anoxic conditions than the other *Methylophaga* sp. The denitrification system at the Montreal Biodome, although contained a deoxygenation reactor, was not fully air-tight and may have sufficient O_2_ (0.3–0.8 mg O_2_ /L; (Labbé *et al*., 2003) for the development of obligate aerobic *Methylophaga* sp. such as *M. frappieri* strain JAM7 that we isolated from this system (Auclair *et al*., 2010).

The relative expression profiles of strain GP59 in the *RR biofilm* were clearly different than those found in (i) the *Original Biofilm*, (ii) the ASW-medium, batch-static biofilm cultures, and (iii) the planktonic cultures with the specific *Methylophaga* medium (*Methylophaga*-1403). These results strongly suggest that the medium, the physiological conditions (planktonic, biofilm) and the operating conditions (anoxic, oxic) have a strong impact on the overall gene expression profiles in strain GP59 (Di Capua *et al*., 2022).

The relative expression profiles of genes involved in N-cycle in strain GP59 have a similar pattern in the four *RR biofilm* samples and in the PlkAN cultures, suggesting that the operating conditions (planktonic cultures versus recirculating biofilm reactor) did not alter the expression profiles of strain GP59 regarding the N-cycle despite changes in C/N ratio in the recirculating reactor. The relative expression patterns of the gene clusters of the *nar1* and *nor1* systems and genes involved in NO response seem to have been more solicitated under these conditions, contrary to the *nar2* and *nor2* systems, which were more involved under the oxic conditions or in the *Original Biofilm*.

The relative expression profiles of the four *xoxF* genes in strain GP59 covariated with the four *RR biofilm* samples. The XoxF-type lanthanide (Ln^3+^)-dependent methanol dehydrogenases are widespread among bacteria. Ln^3+^ is an important co-factor that regulates the expression *xoxF* and *mxa* gene cluster (encoding Ca^2+^-dependent methanol dehydrogenase) (Chistoserdova & Kalyuzhnaya, 2018). We do not know if Ln^3+^ was included in the IO formulation, or whether it exists naturally as trace element. However, the fact that the *xoxF* genes had higher relative transcript levels in the *RR biofilm* samples in opposition of those of *mxa* that was higher in the oxic planktonic cultures, suggest important roles of XoxF of the carbon dynamics in biofilms under anoxic conditions as alternative methanol dehydrogenases (Chistoserdova & Kalyuzhnaya, 2018).

Along with the *xoxF* expression patterns, similar results were observed with 10 out of the 11 riboswitches with higher relative transcript levels in the *RR biofilm* samples. Examination of the genes that flanked the eleven riboswitches showed divers functions. Four of them (Ribo4, 5, 6, 7; c-di-GMP type) are found in close vicinity (18 kb apart), and each flanked a gene encoding protein with PEP-CTERM domain, which seems to be involved in exopolysaccharide-associated protein sorting systems (Haft *et al*., 2006, Villemur *et al*., 2019). Furthermore, genes surrounding these coding sequences encode proteins involved in exopolysaccharide biosynthesis and export. In Gram-negative bacteria, many synthase-dependent exopolysaccharide secretion systems are post-translationally regulated by an inner-membrane c-di-GMP receptor (Whitney & Howell, 2013). Interestingly, Ribo5 covariated with the planktonic anoxic cultures (PlkAN), whereas the three others with the *RR biofilm* samples. These results suggest that specific controls of these pathways that would affect the surface of bacteria are important in the fate of the bacterial physiology (biofilm vs planktonic). It is known that c-di-GMP is a key factor in bacteria to regulate several genes involving in the switch between planktonic and biofilm. One mechanism of biofilm development is the surface attachment, which requires specific adherent organelles such flagella, fimbriae or pili (Martínez & Vadyvaloo, 2014).

## Conclusions

Our results suggest that large-scale denitrification systems would benefit from long term operation with aging microbial community evolving to more diversified populations, which can sustain broad range of C/N changes. This type of microbial community can also adapt more readily to changes in the carbon source (C-1 carbon to complex carbons or mixture of both). For instances, in aquaculture, feeding and fish feces can add diverse carbon sources and generate higher levels of NO_3_^−^ in water, which can disturb the C/N settings in the denitrification system. We believe that knowing the evolution of the biofilm microbial community during the operating conditions is an important outcome of our research. It will certainly help to understand the microbiology of these bioprocesses, to what is hidden behind all these engineering setups.

## Data availability statement

The datasets presented in this study can be found in online repositories. Raw sequencing data were deposited in Sequence Read Archive (SRA) at the National Center for Biotechnology Information (NCBI: https://www.ncbi.nlm.nih.gov/). Metatranscriptomes of Condition 1, Condition 4, Condition 6 and Condition 8: Bioproject PRJNA1194547 and Biosamples SAMN45188687 to SAMN45188690. Original biofilm metatranscriptome: Bioproject PRJNA744510 and Biosamples SAMN20104466. Metatranscriptomes of 300N23C, 900N30C and 0%NaCl: Bioproject PRJNA525230 and Biosamples SAMN11029462, SAMN11029465 and SAMN11029466. Transcriptomes of strain GP59 planktonic cultures (PlkO1 to 3 and PlkON1 to 3): Bioproject PRJNA1072961 and Biosamples SAMN39755719 to SAMN39755725; and PlkAN1 to 3: Bioproject PRJNA525230 and Biosamples SAMN11043171 to SAMN11043173. For the 16S rRNA gene amplicon sequences (D0, Condition 5-1, Condition 5-2 and Condition 6-2): Bioproject PRJNA1194547 and Biosamples SAMN45195935 to SAMN45195938, and for the Original Biofilm: Bioproject PRJNA524642 and Biosamples SAMN11029470.

Assembled contigs from metatranscriptomes performed at Joint Genome Institute IMG/MER, (Condition 1, Condition 4, Condition 6, Condition 8 and Original Biofilm), the GOLD IDs in IMG Database Analysis ID: Ga0510006 to Ga0510010.

## Author contributions

Livie Lestin conceived and designed the experiments, performed the experiments, analyzed the data, prepared figures and/or tables, authored or reviewed drafts of the article, and approved the final draft.

Richard Villemur conceived and designed the experiments, analyzed the data, prepared figures and/or tables, authored or reviewed drafts of the article, and approved the final draft.

## Funding

The authors disclosed the following grant information: Natural Sciences and Engineering Research Council of Canada: # RGPIN-2016-06061.

## Notes

### Competing Interest Statement

The authors have declared no competing interest.

